# Gut physiology of rainbow trout (*Oncorhynchus mykiss*) is influenced more by short-term fasting followed by refeeding than by feeding fishmeal-free diets

**DOI:** 10.1101/2023.05.24.542126

**Authors:** Laura Frohn, Diogo Peixoto, Frédéric Terrier, Benjamin Costas, Jérôme Bugeon, Christel Cartier, Nadège Richard, Karine Pinel, Sandrine Skiba-Cassy

## Abstract

Supplementing a fishmeal-free diet with yeast extract improves rainbow trout (*Oncorhynchus mykiss*) growth performance and modulates the hepatic and intestinal transcriptomic response. These effects are often observed in the long term, but are not well documented after short periods of fasting. Fasting for a few days is a common practice in fish farming, especially before handling the fish, such as for short sorting, tank transfers and vaccinations. In the present study, rainbow trout were subjected to a four-day fast and then refed, for eight days, a conventional diet containing fishmeal (control diet) or alternative diets composed of terrestrial animal by-products supplemented or not with a yeast extract. During the refeeding period alone, most of the parameters considered did not differ significantly in response to the different feeds. Only the expression of claudin-15 was upregulated in fish fed the yeast-supplemented diet compared to the control diet. Conversely, fasting followed by refeeding significantly influenced most of the parameters analyzed. In the proximal intestine, the surface area of villi significantly increased and the density of goblet cell tended to decrease during refeeding. Although no distinct plasma immune response or major signs of gut inflammation were observed, some genes involved in the structure, complement pathway, antiviral functions, coagulation and endoplasmic reticulum stress response of the liver and intestine were significantly regulated by refeeding after fasting. These results indicate that short-term fasting, as commonly practiced in fish farming, significantly alters the physiology of the liver and intestine regardless of the composition of the diet.

## 1. Introduction

Aquaculture rearing practices and feeds changed greatly during the 20^th^ century. Development of extruded pellets in the late 1980s and the gradual replacement of fishmeal and fish oil with plant ingredients (e.g. whole grains, oilseeds, legumes) helped address the constraints faced by the sector, particularly in reducing human pressures on wild forage-fish populations (1, 2). From 1990-2020, especially for Atlantic salmon (*Salmo salar*), the contents of fish oil and fishmeal in feed decreased from 65.4% and 24.0%, respectively, to 12.1% and 10.3%, respectively, with no influence on zootechnical performances (3). Nonetheless, most carnivorous fish species, such as salmonids (salmon and trout), are sensitive to dietary changes, which prohibits completely replacing fishmeal and fish oil in their feed. Thus, for these species, plant-based diets are likely to influence feed intake, nutrient digestion, absorption and assimilation (e.g. metabolism) (4–6). These diets can also degrade intestinal health by causing inflammation in the digestive tract, likely due to the presence of anti-nutritional factors (e.g. oligosaccharides, lectins, saponins) in the plant ingredients (7, 8). Therefore, plant-based diets are likely to decrease the growth and degrade the health of the fish, thus rendering them more sensitive to stress (9, 10).

In addition to plant ingredients, the aquaculture sector is considering other replacements for marine ingredients, such as insect meals, microalgae and industrial by-products from terrestrial animals. However, their use is restricted due to variable growth performances that result from their inconsistent quality and nutritional values, as well as the amounts included in diets (11). Including functional ingredients can meet the objectives of maintaining acceptable growth performances and mitigating adverse effects of alternative diets. For example, yeast can be included whole or in derived form (i.e. cell wall, cytosolic fraction, purified components). Including yeast in feed has been shown to provide many benefits for most terrestrial and aquatic livestock, such as improvements in feed palatability, nutrient digestibility (due to enzyme production and intestinal pH regulation), the immune system and pathogen resistance (due to bioactive components such as MOS, β-glucan, nucleotides and peptides), the quality of meat and milk, egg production of poultry and growth performances of fish and pigs (12–14).

The performance of new feeds or the benefits of feed additives are often examined over the long term to assess effects on growth without necessarily analyzing the early physiological responses of animals. However, a study of Atlantic salmon fed a soybean meal diet showed that diet can have effects on the first day of exposure, particularly on intestinal histopathology and on the regulation of genes involved in immunity, detoxification, cellular repair and metabolism (15). These early effects should be reconsidered, since periods of fasting and refeeding are frequently implemented in aquaculture. On fish farms, short-term fasting is strongly recommended and commonly applied during periods of stress (i.e. animal handling, transport between different production sites, epizootic situations) or in order to induce a compensatory growth (16–18). It lowers the metabolic rate of fish and causes the energy required for digestion to be directed to vital functions such as maintaining cellular homeostasis and immune functions, without necessarily causing weight loss. Like birds and mammals, fish in their natural environment are frequently subjected to periods of fasting during winter, migration periods and reproduction events (19, 20). Even when short, these periods influence gut microbiota (21), metabolism (22), and digestive and hepatic physiology (20, 23, 24).

In this study, we evaluated the influences of a short fasting period followed by refeeding on the histology of the intestine and parameters related to defense and protection mechanisms. In this context, we compared three diets: a conventional efficient diet containing fishmeal and fish oil, an alternative diet based on plant ingredients and terrestrial animal by-products (which had decreased the growth performance of juvenile rainbow trout (*Oncorhynchus mykiss*) in a previous study (25)) and the same diet supplemented with yeast extract (which had significantly increased their growth).

## 2. Materials and methods

### Ethical statement

Rainbow trout were reared at the French National Research Institute for Agriculture, Feed and Environment (INRAE) experimental facility (26), authorized for animal experimentation by the French Veterinary Service (A64-495-1 and A40-228-1). The technical staff received training and authorization to conduct experiments. The experiment was conducted in strict agreement with Directive 2010/63/EU of European Union regulations on the protection of animals used in scientific research, and the National Guidelines for Animal Care of the French Ministry of Research (decree no. 2013-118, 02 Jan 2013). The experiment did not need approval from the ethical committee (C2EA-73) since it complied with current rearing standards.

### Feed formulation and experimental design

Rainbow trout, with mean body weight of 232 ± 39 g, were reared in 80 L tanks supplied continuously with well-oxygenated freshwater at 17 °C. Three experimental feeds were formulated to meet the nutritional requirements of the rainbow trout according to NRC recommendations (27). The control diet (CTL) was formulated as close as possible to a commercial feed and contained 19% fishmeal and 7.0% of fish oil. Two processed animal protein diets (10% dehydrated poultry protein, 6% hydrolyzed feather meal and 1 % poultry and pig blood meal) were also formulated: one not supplemented (PAP) and one supplemented with yeast extract (PAP+YE). The yeast extract used contained the cytosolic fraction of the yeast *Saccharomyces cerevisiae* (Prosaf®, Phileo by Lesaffre, Marcq-en-Barœul, France). The contents of other ingredients in the experimental feeds varied (Table 1).

**Table 1.**
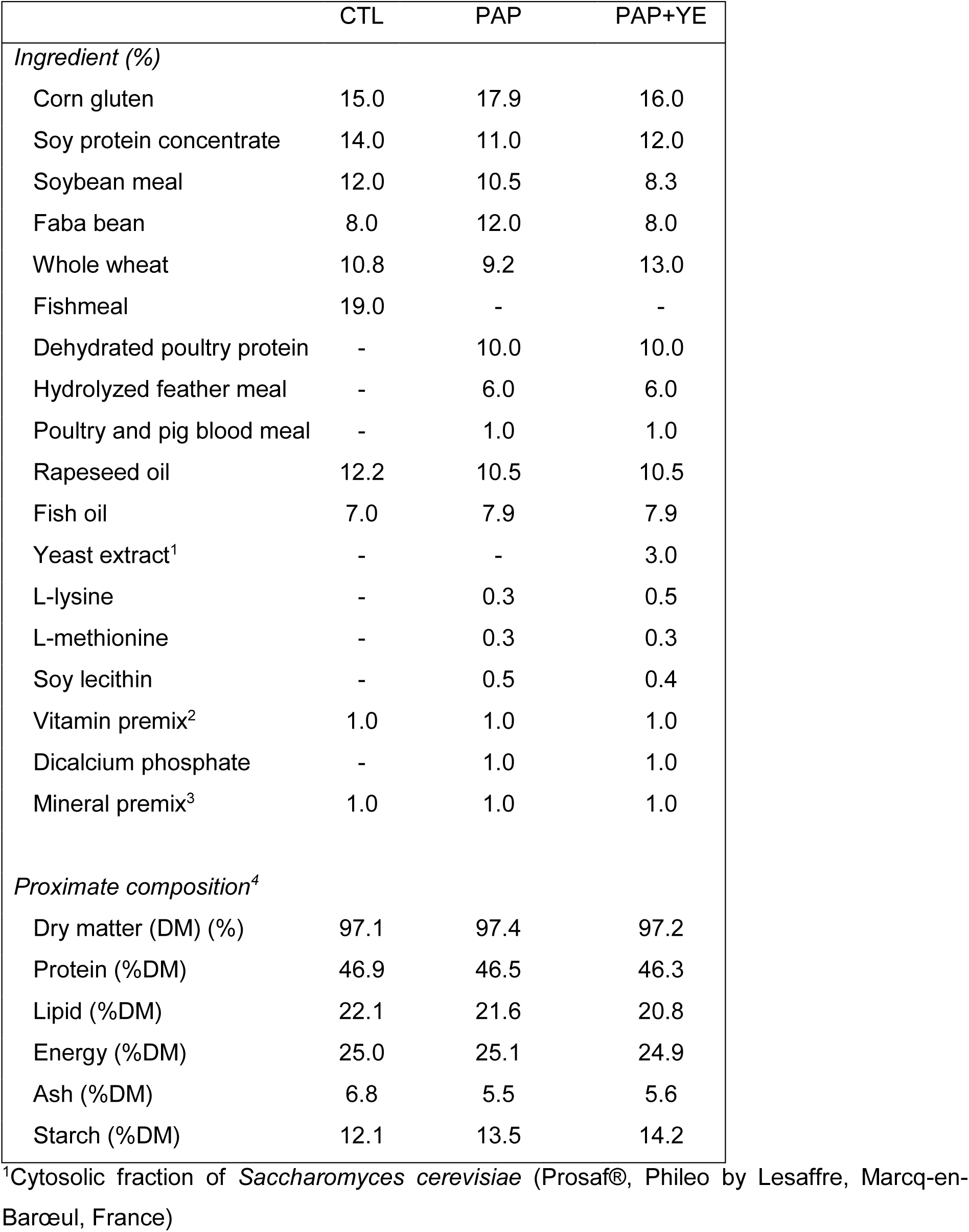
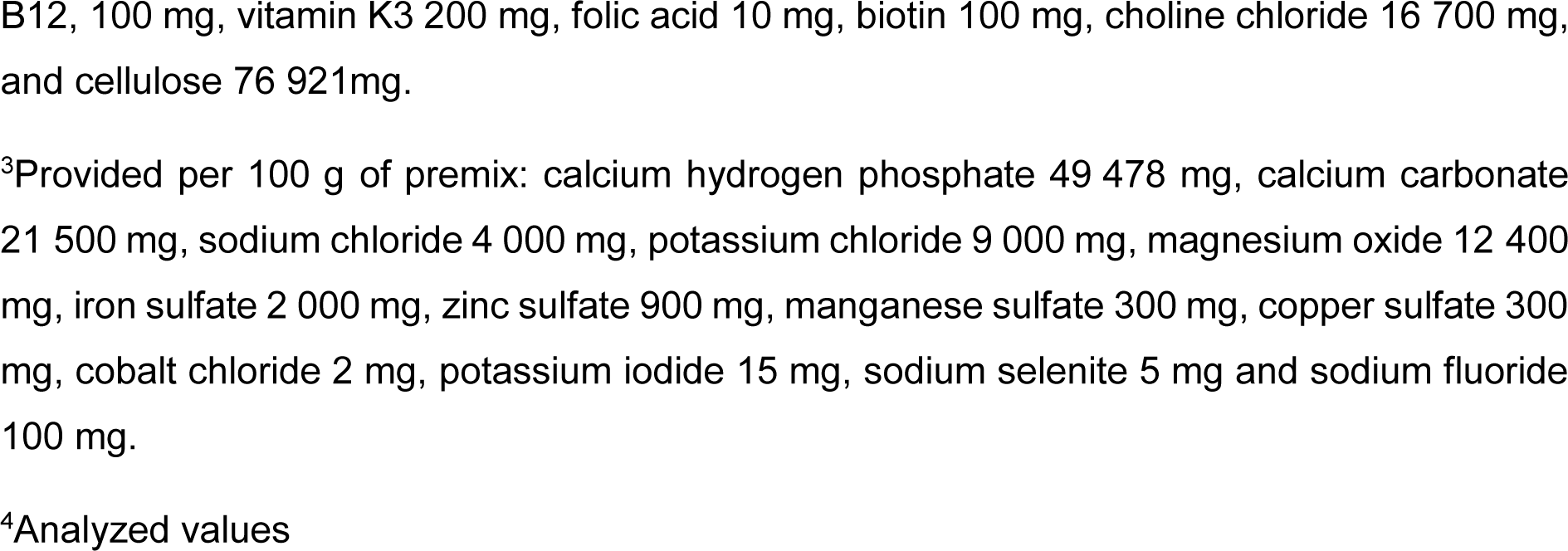
Ingredients and proximate composition of three experimental diets: commercial-like feed (CTL), processed animal protein feed (PAP) and PAP with 3% yeast extract (PAP+YE).

|  | CTL | PAP | PAP+YE |
| --- | --- | --- | --- |
| *Ingredient (%)* |  |  |  |
| Corn gluten | 15.0 | 17.9 | 16.0 |
| Soy protein concentrate | 14.0 | 11.0 | 12.0 |
| Soybean meal | 12.0 | 10.5 | 8.3 |
| Faba bean | 8.0 | 12.0 | 8.0 |
| Whole wheat | 10.8 | 9.2 | 13.0 |
| Fishmeal | 19.0 | - | - |
| Dehydrated poultry protein | - | 10.0 | 10.0 |
| Hydrolyzed feather meal | - | 6.0 | 6.0 |
| Poultry and pig blood meal | - | 1.0 | 1.0 |
| Rapeseed oil | 12.2 | 10.5 | 10.5 |
| Fish oil | 7.0 | 7.9 | 7.9 |
| Yeast extract<sup>1</sup> | - | - | 3.0 |
| L-lysine | - | 0.3 | 0.5 |
| L-methionine | - | 0.3 | 0.3 |
| Soy lecithin | - | 0.5 | 0.4 |
| Vitamin premix<sup>2</sup> | 1.0 | 1.0 | 1.0 |
| Dicalcium phosphate | - | 1.0 | 1.0 |
| Mineral premix<sup>3</sup> | 1.0 | 1.0 | 1.0 |
| *Proximate composition<sup>4</sup>* |  |  |  |
| Dry matter (DM) (%) | 97.1 | 97.4 | 97.2 |
| Protein (%DM) | 46.9 | 46.5 | 46.3 |
| Lipid (%DM) | 22.1 | 21.6 | 20.8 |
| Energy (%DM) | 25.0 | 25.1 | 24.9 |
| Ash (%DM) | 6.8 | 5.5 | 5.6 |
| Starch (%DM) | 12.1 | 13.5 | 14.2 |

<sup>1</sup>Cytosolic fraction of *Saccharomyces cerevisiae* (Prosaf®, Phileo by Lesaffre, Marcq-en-Barœul, France)
<sup>2</sup>Provided per 100 g of premix: vitamin A 500 000 IU, vitamin D3 250 000 IU, vitamin E 5 00 mg, vitamin C 1 429 mg, vitamin B1 10 mg, vitamin B2 50 mg, vitamin B3 100 mg, vitamin B5 200 mg, vitamin B6 30 mg, vitamin B7 3 000 mg, vitamin B8 100 mg, vitamin B9 10 mg, vitamin
B12, 100 mg, vitamin K3 200 mg, folic acid 10 mg, biotin 100 mg, choline chloride 16 700 mg, and cellulose 76 921mg.
<sup>3</sup>Provided per 100 g of premix: calcium hydrogen phosphate 49 478 mg, calcium carbonate 21 500 mg, sodium chloride 4 000 mg, potassium chloride 9 000 mg, magnesium oxide 12 400 mg, iron sulfate 2 000 mg, zinc sulfate 900 mg, manganese sulfate 300 mg, copper sulfate 300 mg, cobalt chloride 2 mg, potassium iodide 15 mg, sodium selenite 5 mg and sodium fluoride 100 mg.
<sup>4</sup>Analyzed values

Rainbow trout were distributed among four tanks, which contained 30 fish each. After batching, the fish were acclimated and fed a commercial feed (T3P Omega®, Skretting, Fontaine-les-Vervins, France) for three days (distributed daily to visual satiation) and then fasted for four days to empty their digestive tracts completely. Then, each experimental diet was randomly assigned to three tanks, and fish were fed twice a day until visual satiation, for eight days. The fish in the fourth tank, which were not refed after fasting, were used to represent day 0 of the experiment, when six of them were sampled by anesthetizing and euthanizing them in successive baths of tricaine (50 mg/L and 150 mg/L, respectively). On days 2, 5 and 8, fish in each of the other tanks were sampled in the same way, 6 hours after the last meal. Samples of the liver and distal intestine were collected, soaked in RNA later and immediately immersed in liquid nitrogen and stored at -80 °C until molecular analyses was performed. Additional samples of the proximal and distal intestine were fixed in a 10% buffered formalin solution before being processed for histological analysis. Blood was drawn from the caudal vein using an EDTA-treated syringe, centrifuged at 3000 *g* for 10 minutes and the plasma extracted was stored at -20°C until plasma immune markers were quantified.

### Plasma immune markers

Activities of the alternative complement pathway, lysozyme and peroxidase were measured in the plasma, as described by Frohn *et al.* (25). Total antiprotease activity was determined by assessing the ability of plasma to inhibit trypsin activity, as described by Peixoto *et al.* (28).

### Histology of intestine

According to recommendations of Feldman & Wolfe (29), samples of proximal and distal intestine were fixed in buffered formalin, dehydrated in successive ethanol baths, clarified in xylene and embedded in paraffin blocks. Transverse intestine sections 2 µm thick were cut using a semi-automatic rotating microtome (HM 340E, Microm Microtech, Brignais, France). Tissue was then stained with periodic acid-Schiff alcian blue to assess gut morphology and count the number of goblet cells in the intestinal mucosa, as it has been described by Frohn *et al.* (25).

### Image analysis

Images of the intestinal mucosa were obtained using a microscope (DMRB, Leica, Wetzlar, Germany) equipped with a digital camera (DP71 1.4M pixels, Olympus, Tokyo, Japan). The images were then processed using two macros that were created using the MorphoLibJ (30) and StarDist (31) plugins of FIJI 1.53t software (32). The first macro measured the surface area (µm²) of a given villus and counted the number of goblet cells in it, in order to calculate the density of goblet cells (per mm²). The cells were detected using the StarDist Versatile (H&E nuclei) model, and each villus of interest was manually delineated (polygon-selection tool). The cells inside the villus were manually corrected if incorrectly segmented (Fig. 1). The second macro measured villi height (µm) manually using the segmented-line tool of FIJI. Five villi per section were analyzed.

**Figure 1.**
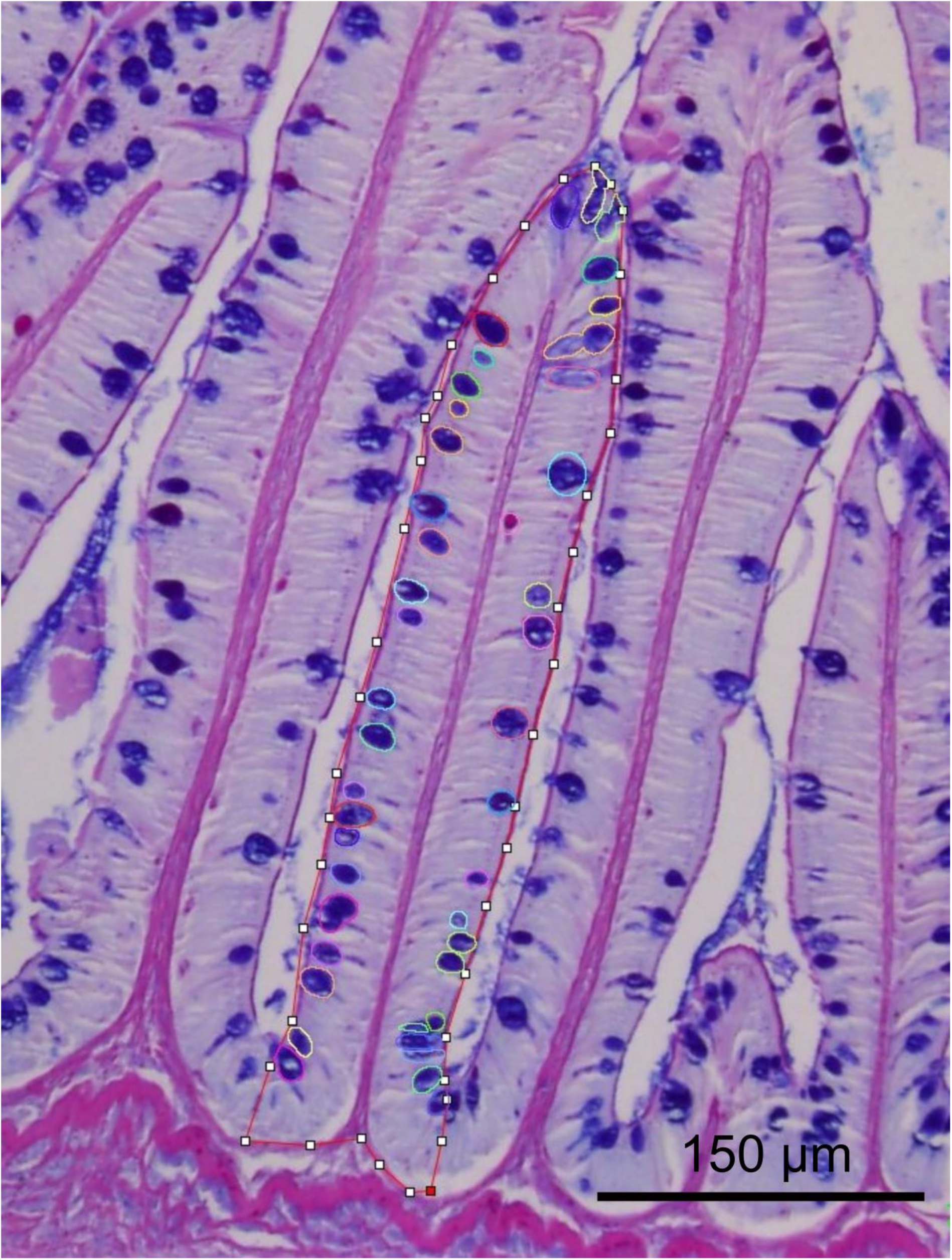
Example of manual delineation of a villus (red line) and detection of goblet cells (circles) using FIJI software.

### Molecular analysis: RNA extraction and real-time RT-qPCR

Total RNA was extracted from the liver, proximal intestine and distal intestine (n = 6 *per* sampling day and diet) according to the miRNeasy Tissue/Cells Advanced Mini Kit protocol (Qiagen, Hilden, Germany). RNA was treated with the Turbo DNA-free kit (Invitrogen, Waltham, Massachusetts, USA) to avoid genomic DNA contamination. RNA integrity was verified on 1% agarose gel, and RNA concentrations were quantified using a spectrophotometer (Nanodrop® ND1000, Thermo Scientific, Waltham, Massachusetts, USA). The cDNA synthesis was performed with 750 ng of total RNA using the SuperScript III reverse transcriptase (Invitrogen, USA) and random primers (Promega, Madison, Wisconsin, USA) according to manufacturers’ instructions. Then, for each gene of interest, real-time RT-qPCR was performed using the Lightcycler® 480 II system (Roche Diagnostics, Basel, Switzerland) with primers designed based on (i) genes identified in a previous study of Frohn et al. (25) using the same diet during a long-term feeding period and (ii) supplemental genes that are key markers of the pathways of interest (Table 2). These genes were involved in immunity and inflammation, structure, coagulation functions, cell protection, antiviral functions and endoplasmic reticulum stress.

**Table 2.**
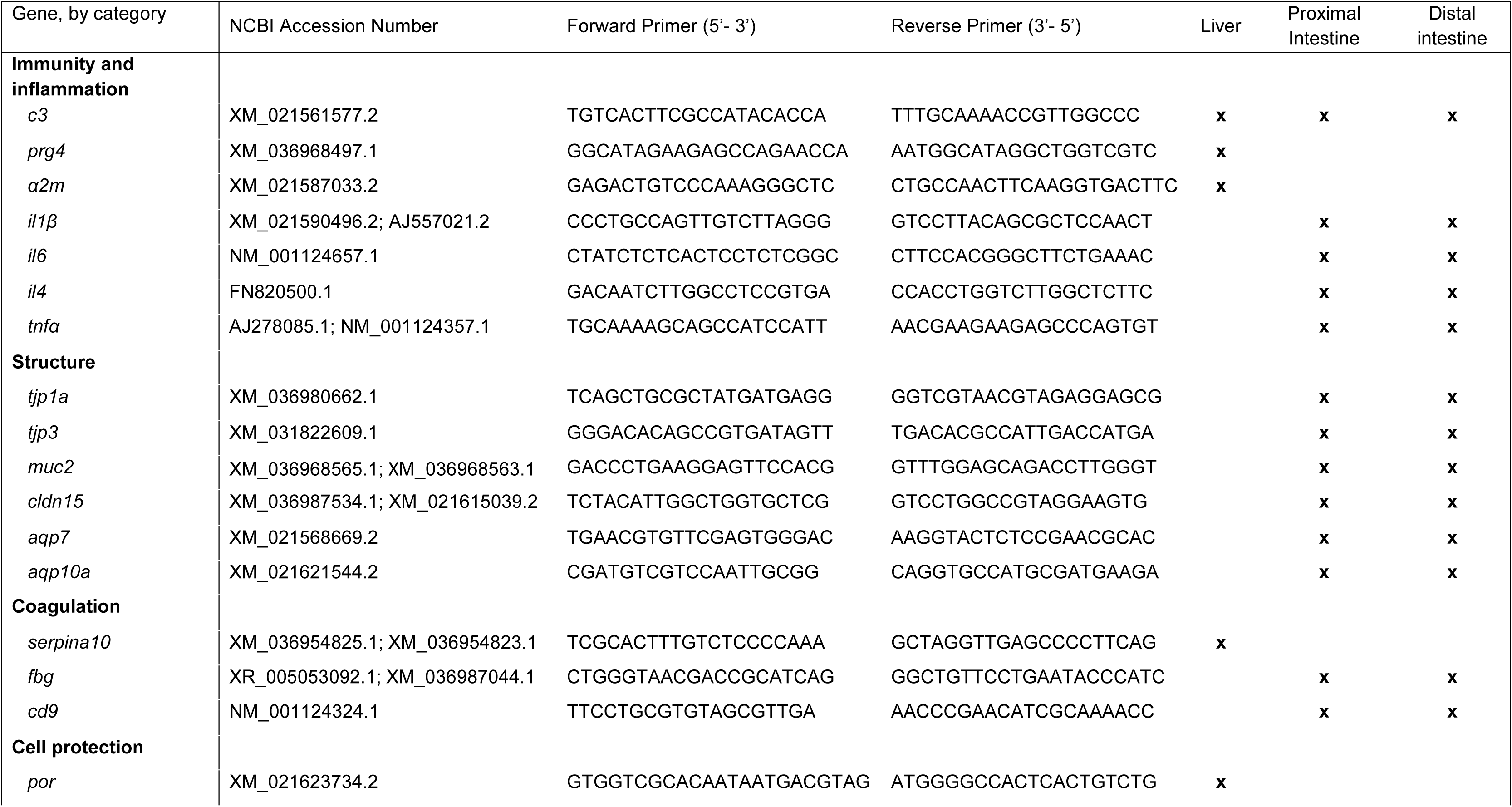
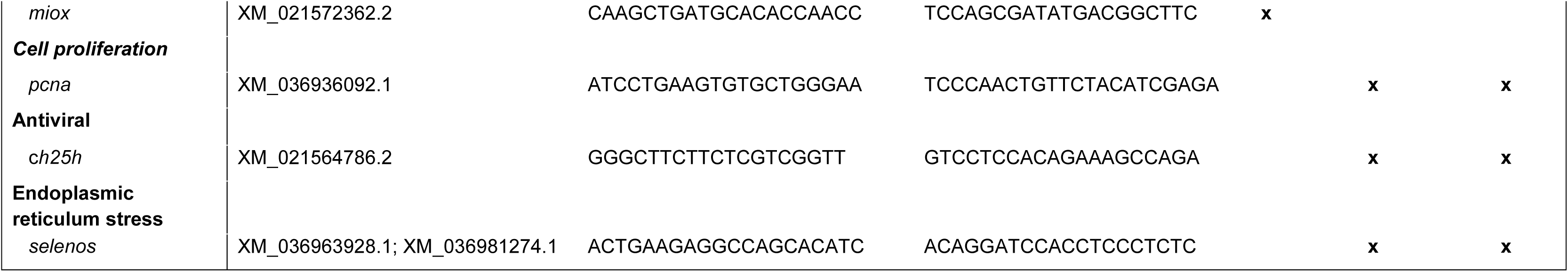
Primer sequences for RT-qPCR analysis of the liver and proximal and distal intestine

Real-time RT-qPCR was performed using a reaction mix containing 2 µL of diluted cDNA, 3 µL of Light Cycler 480 SYBR® Green I Master mix (Roche Diagnostics, Switzerland), 0.24 µL of each primer (10X) and 0.52 µL of RNase- and DNase-free water (Thermo Fisher Scientific, USA). Negative controls constituted of RT- and cDNA-free samples. Each RT-qPCR assay was deposited in triplicates on a FrameStar® 384-well skirted qPCR plate (Roche Diagnostics, Switzerland). The qPCR program was initiated at 95°C for 10 minutes to denature the cDNA and activate the TAQ polymerase enzyme in a thermocycler (Lightcycler® 480 II Roche thermocycler, Roche Diagnostics, Switzerland). The initiation was followed by 45 amplification cycles, each consisting of successive thermal steps (15 sec at 95°C, 10 sec at 60°C and 15 sec at 72 °C). Melting curves (0.5°C/10 sec from 65 to 95°C) were run at the end of each amplification cycle to confirm the specificity of the reaction. The relative expression of genes was quantified using the ΔΔCT method (33). Expression levels were normalized relative to those of the fasted fish using elongation factor EF1α (forward 5’-TCCTCTTGGTCGTTTCGCTG-3’; reverse 3’-ACCCGAGGGACATCCTGTG-5’), 18S rRNA (forward 5’-CGGAGGTTCGAAGACGATCA-3’; reverse 3’-TCGCTAGTTGGCATCGTTTAT-5’) and β-actin (forward 5’-GATGGGCCAGAAAGACAGCTA-3’; reverse 3’-TCGTCCCCAGTTGGTGACGAT-5’) for the liver, proximal intestine and distal intestine, respectively.

### Statistical analysis

Plasma immune markers, histological parameters and gene expressions were statistically analyzed using the Rcmdr package of R software (version 4.1.2). For each dataset, normality of the distributions and homogeneity of the variances were assessed using a Shapiro-Wilk’s test and a Levene’s test, respectively. When these prerequisites were met, the effects of the diet and duration of refeeding (days 2, 5 and 8) were analyzed using two-way analysis of variance (ANOVA) by excluding the fasted fish. Then, the influence of time (i.e. sampling day) was tested independent of the diet by analyzing the datasets from day 0 (fasted fish) to day 8 using one-way ANOVA. Differences were considered statistically significant at the 5 % threshold.

## 3. Results

### 3.1. Plasma immune markers

Neither time nor diet (one-way ANOVA, p > 0.05) significantly influenced the plasma activities of the alternative complement pathway, lysozyme, antiprotease or peroxidase (two-way ANOVA, p > 0.05) (Fig. 2). However, an interaction between diet and time was observed for plasma lysozyme activity (p < 0.05), perhaps because it decreased significantly in fish fed the PAP diet on day 8 compared to day 2, and compared to the fish fed the CTL diet on days 5 and 8.

**Figure 2.**
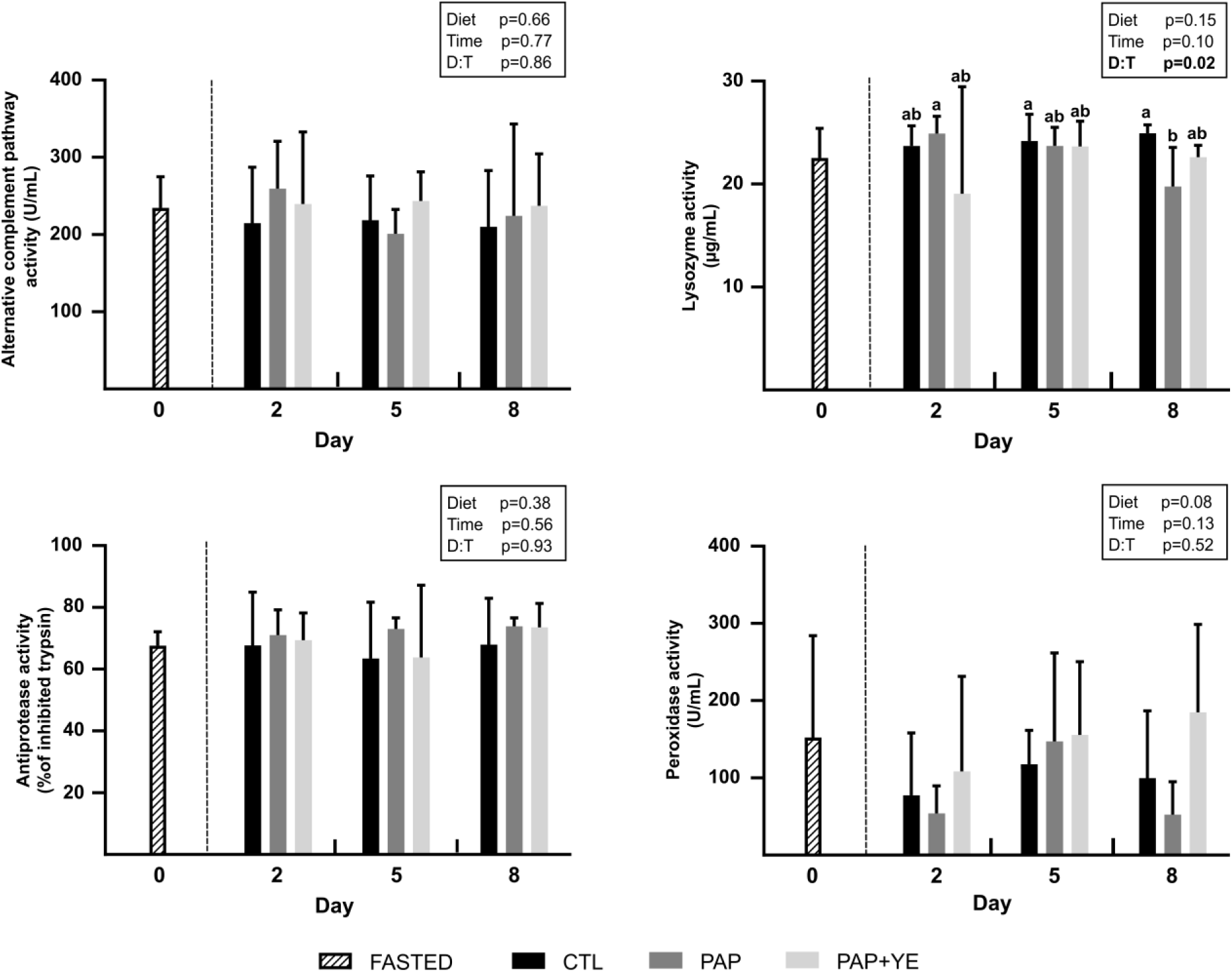
Mean activity (+ standard deviation) (n = 6) of plasma immune markers in rainbow trout after fasting and on three of the eight subsequent days of refeeding. CTL, commercial-like feed; PAP, processed animal protein feed; PAP+YE, PAP with 3% yeast extract. Insets indicate the significance of effects of diet, time and their interaction (D:T). Lowercase letters indicate the significant differences in the interaction between diet and time.

### 3.2. Histological parameters of the proximal and distal intestine

In the proximal and distal intestine, neither diet nor time significantly influenced villi height on days 2, 5 or 8 (two-way ANOVA, p > 0.05), and villi height did not differ significantly between fasted fish (day 0) and refed fish (one-way ANOVA, p > 0.05) (Fig. 3). In the proximal intestine, the diet did not influence villi surface area (two-way ANOVA, p > 0.05), but time did with a significant increase in villi surface area on day 8 compared to those on days 2 and 0 (two-way ANOVA, p < 0.05). The surface area of villi for fasted fish (day 0) and refed fish on day 2 was significantly smaller than that of refed fish on day 8 (one-way ANOVA, p < 0.05). In the distal intestine, neither diet nor time influenced villi surface area on days 2, 5 or 8 (two-way ANOVA, p > 0.05), nor did (one-way ANOVA, p > 0.05).

**Figure 3.**
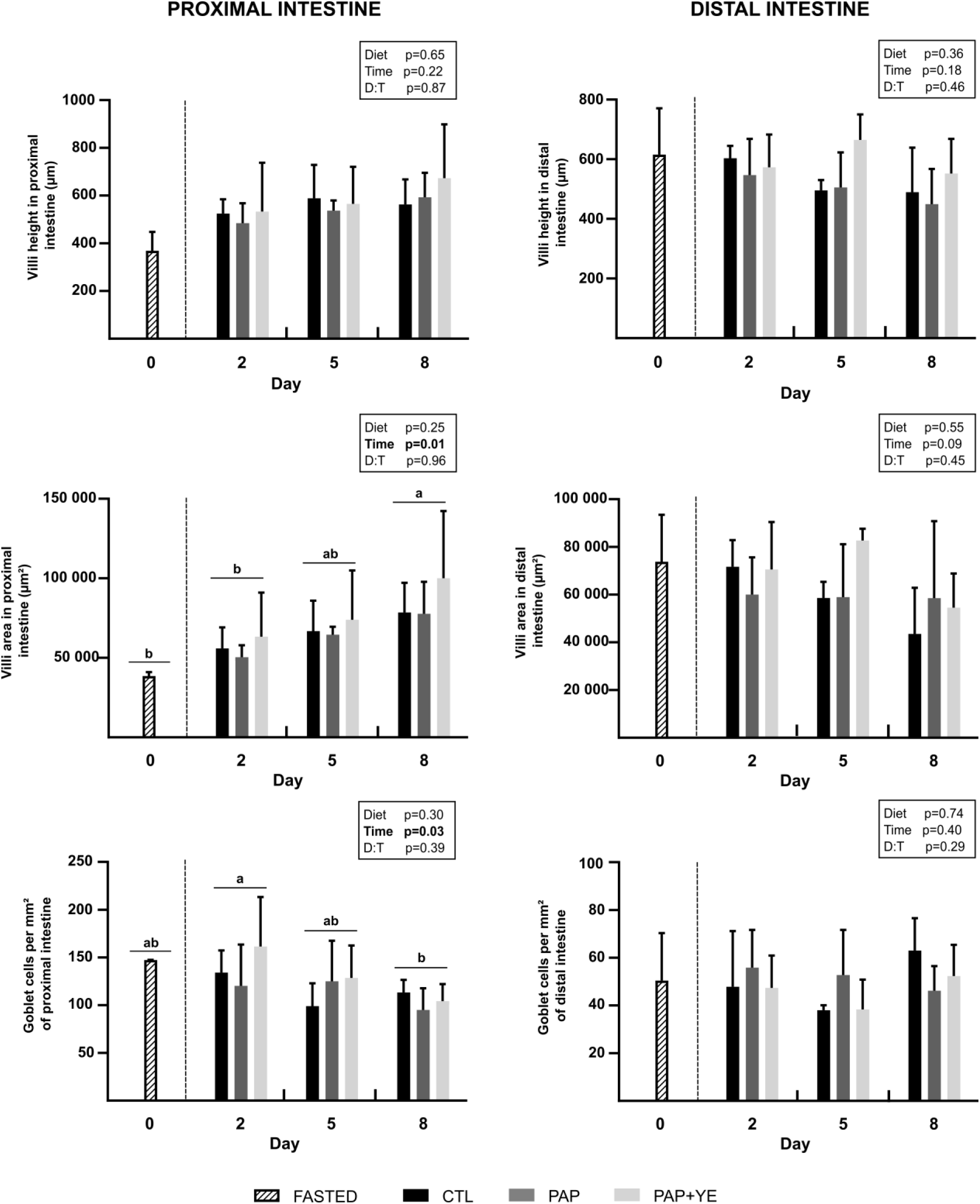
Mean morphometric markers (+ standard deviation) (n = 6) of the proximal and distal intestine of rainbow trout after fasting and on three of the eight subsequent days of refeeding. CTL, commercial-like feed; PAP, processed animal protein feed; PAP+YE, PAP with 3% yeast extract. Insets indicate the significance of effects of diet, time and their interaction (D:T). Lowercase letters indicate significant differences between sampling days.

The diet did not influence the mean density of goblet cells in the proximal or distal intestine (two-way ANOVA, p > 0.05), but refeeding did (two-way ANOVA, p < 0.05), since it was significantly lower on day 8 compared to day 2. Fasted fish (day 0) and refed fish on day 5 had intermediate densities (one-way ANOVA, p < 0.05).

### 3.4. Gene expression in the liver and the proximal and distal intestine

In the liver, the diets did not significantly influence the expression of genes related to inflammation and immunity (i.e. main complement molecule C3 (*c3*); proteoglycan 4 (*prg4*) and alpha-2-macroglobulin isoform X2 (*α2m*)) and to cell protection (i.e. myo-inositol oxygenase (*miox*)) (two-way ANOVA, p > 0.05). However, their expression significantly decreased during refeeding (two-way ANOVA, p <0.05) (Fig. 4). Expression of the *c3* and *prg4* genes was significantly downregulated on day 8 compared to that on day 2 and was intermediate on days 0 and 5. Expression of the acute-phase protein encoded by *a2m* was significantly upregulated in fasted fish (day 0) compared to that on day 8 and was intermediate on days 2 and 5. Expression of the *miox* gene peaked on days 0 and 5, was intermediate on day 2, and was significantly downregulated on day 8. Expression of the *por* (NADPH-cytochrome P450 reductase) and *serpina10* (protein Z-dependent protease inhibitor-like) genes, involved in cell protection and coagulation respectively, was to too low in the liver to be quantified adequately (i.e. ≥ 32 amplification cycles (Ct)).

**Figure 4.**
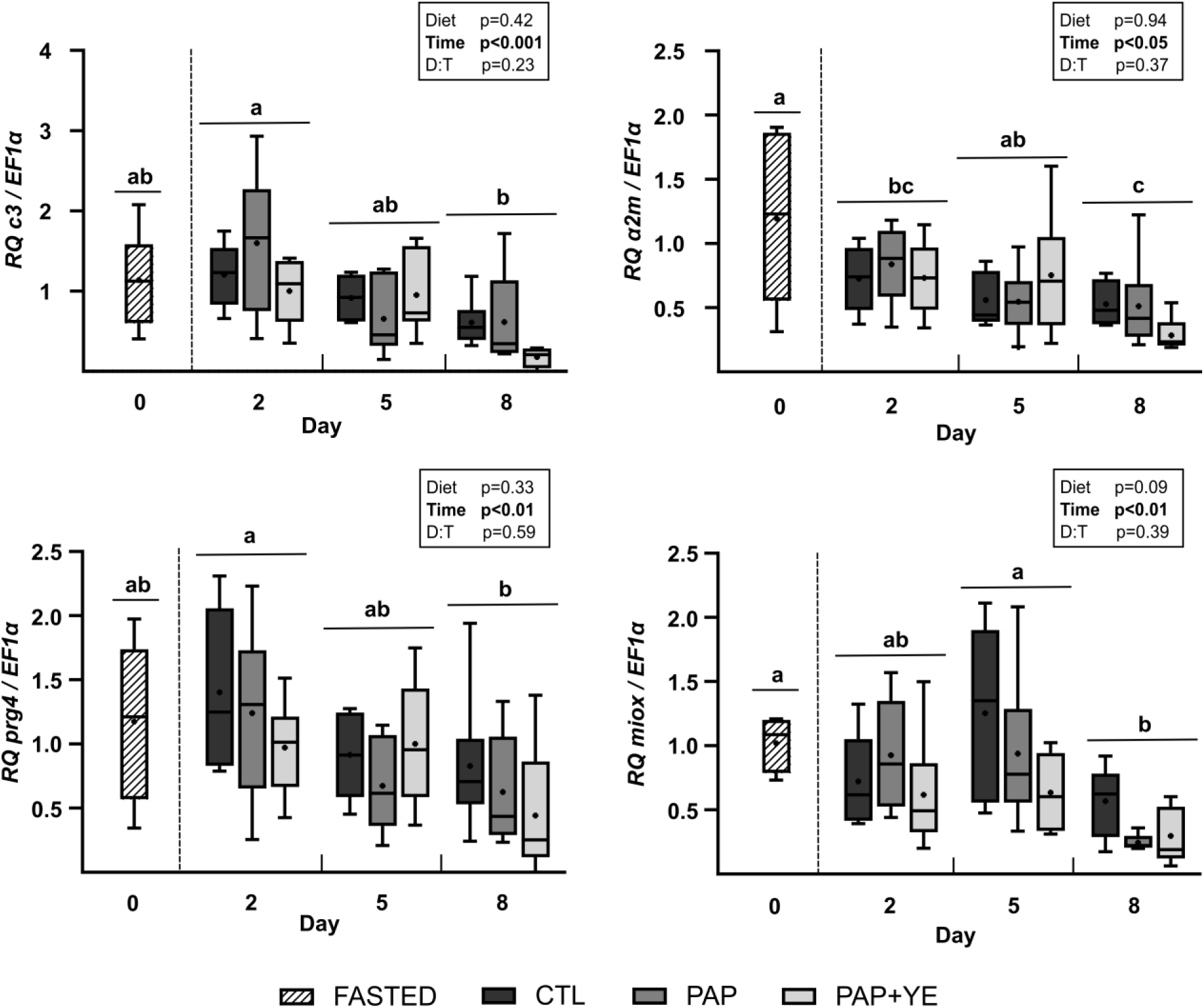
Mean normalized expression (+ standard deviation) (n = 6) of genes related to immunity, inflammation and cell protection in the liver of rainbow trout after fasting and on three of the eight subsequent days of refeeding. CTL, commercial-like feed; PAP, processed animal protein feed; PAP+YE, PAP with 3% yeast extract. *c3*, main complement molecule C3; *prg4*, proteoglycan 4; *miox*, myo-inositol oxygenase; *α2m*, alpha-2-macroglobulin isoform X2. Insets indicate the significance of effects of diet, time and their interaction (D:T). Lowercase letters indicate significant differences between sampling days.

In the proximal intestine, expression of genes related to immunity and inflammation (i.e. *c3*, interleukin-1 beta (*il1β*), interleukin-6 (*il6*), interleukin-4 (*il4*) and tumor necrosis factor alpha (*tnfα*)) was too low (Ct ≥ 32) to be quantified adequately using RT-qPCR. In the distal intestine, the expressions levels of genes *c3*, apolipoprotein A-II (*apoa2)*, *il4*, *il6* and *tnfα* were also too low (Ct > 32) to be analyzed. Interleukin *il1β* was the only gene related to inflammation that was sufficiently expressed in the distal intestine, in which its expression was related to time during refeeding (two-way ANOVA, p<0.001). Compared to fasted fish, the expression of *il1β* increased after 2 days of refeeding, and then returned to the fasted level on days 5 and 8 (one-way ANOVA, p < 0.001). Its expression was not influenced by the diet (two-way ANOVA, p > 0.05), but was significantly influenced by the interaction between diet and time due to upregulation of *il1β* on day 2 in fish fed the PAP compared to fish fed the CTL diet on day 5 (two-way-ANOVA, p < 0.05). Expression of the cholesterol 25-hydroxylase gene (*ch25h*), involved in anti-viral defense, was too low (Ct ≥ 32) in the proximal intestine to be quantified adequately, but was significantly down regulated in the distal intestine of fish on day 5 compared to fasted fish (one-way ANOVA, p < 0.05). However, diet, time and their interaction had no significant effect (two-way ANOVA, p > 0.05) (Fig. 5).

**Figure 5.**
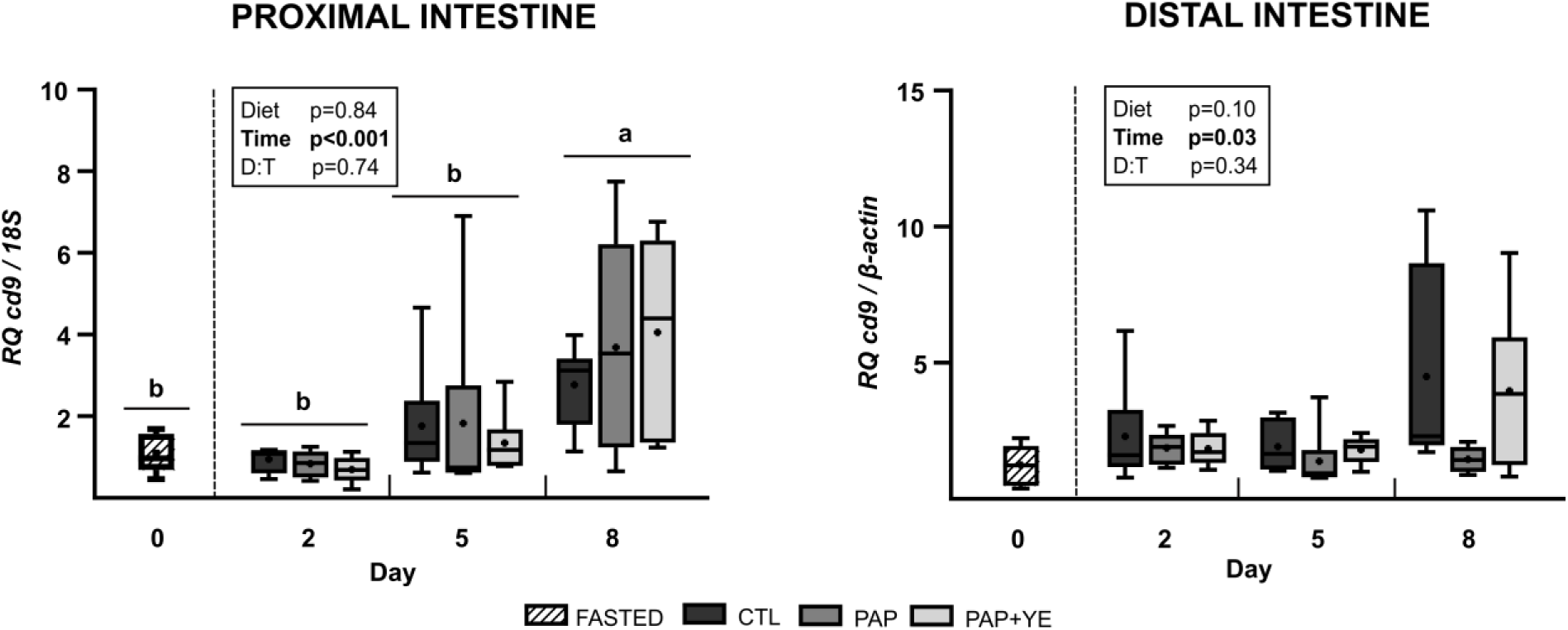
Mean normalized expression (+ standard deviation) (n = 6) of genes related to inflammation and antiviral functions in the distal intestine of rainbow trout after fasting and on three of the eight subsequent days of refeeding. CTL, commercial-like feed; PAP, processed animal protein feed; PAP+YE, PAP with 3% yeast extract; *il1β*, interleukin-1 beta; *ch25h*, cholesterol 25-hydroxylase. Insets indicate the significance of effects of diet, time and their interaction (D:T). Lowercase letters indicate significant differences between sampling days.

Among the tight-junction proteins analyzed in the proximal and distal part of the intestine, only tight-junction protein-3 (*tjp3*) and claudin-15 (*cldn15*) were expressed (Fig. 6). In both sections of the intestine, expression of tight junction protein-1 (*tjp1*) was too low to be quantified adequately using RT-qPCR (Ct ≥ 32). In the proximal intestine, *tjp3* and *cldn15* were significantly influenced by time (two-way ANOVA, p <0.001) but not by the diet (two-way ANOVA, p > 0.05) during refeeding. Moreover, a significant interaction was observed between diet and time for *tjp3* and *cldn15*, which was related to overexpression of both genes on day 8 in fish fed the PAP diet. Over the entire experiment, time had a significant influence (one-way ANOVA, p < 0.01): both genes had significantly higher expression on day 8 when compared to days 0, 2 and 5 for *tjp3*, and compared to day 2 for *cldn15*. In the distal part of the intestine, *tjp3* expression did not differ during refeeding (two-way ANOVA, p > 0.05), but it was significantly lower in fasted fish, especially compared to refed fish on days 2 or 8 (one-way ANOVA, p < 0.05). Expression of *cldn15* also gradually increased from days 0 to 8, being triggered by refeeding (two-way ANOVA, p < 0.05). Expression of *muc2*, which encodes a protein involved in secretion of intestinal mucus, was upregulated in the proximal and distal intestine on 8 compared to those on days 0, 2 and 5 (one-way ANOVA, p < 0.05 and one-way ANOVA, p < 0.01, respectively). Expression of *muc2* was not influenced by the diets in either section of the intestine (two-way ANOVA, p > 0.05). Expression of the aquaporins *aqp7* and *aqp10a* was too low in both sections of the intestine to be quantified adequately (Ct ≥ 32). Expression of the proliferating cell nuclear antigen (*pcna*) gene, involved in DNA replication and cell proliferation, was regulated by the diet in the proximal intestine, being lower in fish fed the PAP+YE diet than in fish fed the PAP diet (two-way ANOVA, p < 0.05). Its expression was also influenced by refeeding (two-way ANOVA, p < 0.01) and its duration (one-way ANOVA, p < 0.001). Fasted fish had the lowest expression of *pcna* and refeeding induced a gradual increase in *pcna* expression on days 2 and 8, with intermediated expression on day 5. In the distal intestine, time influenced *pcna* expression between fasted and refed fish (one-way ANOVA, p < 0.01). As in the proximal intestine, expression of *pcna* in the distal intestine was lower in fasted fish, increased significantly in refed fish on day 2 and decreased on days 5 and 8, but remained significantly higher than that in fasted fish (Fig. 6).

**Figure 6.**
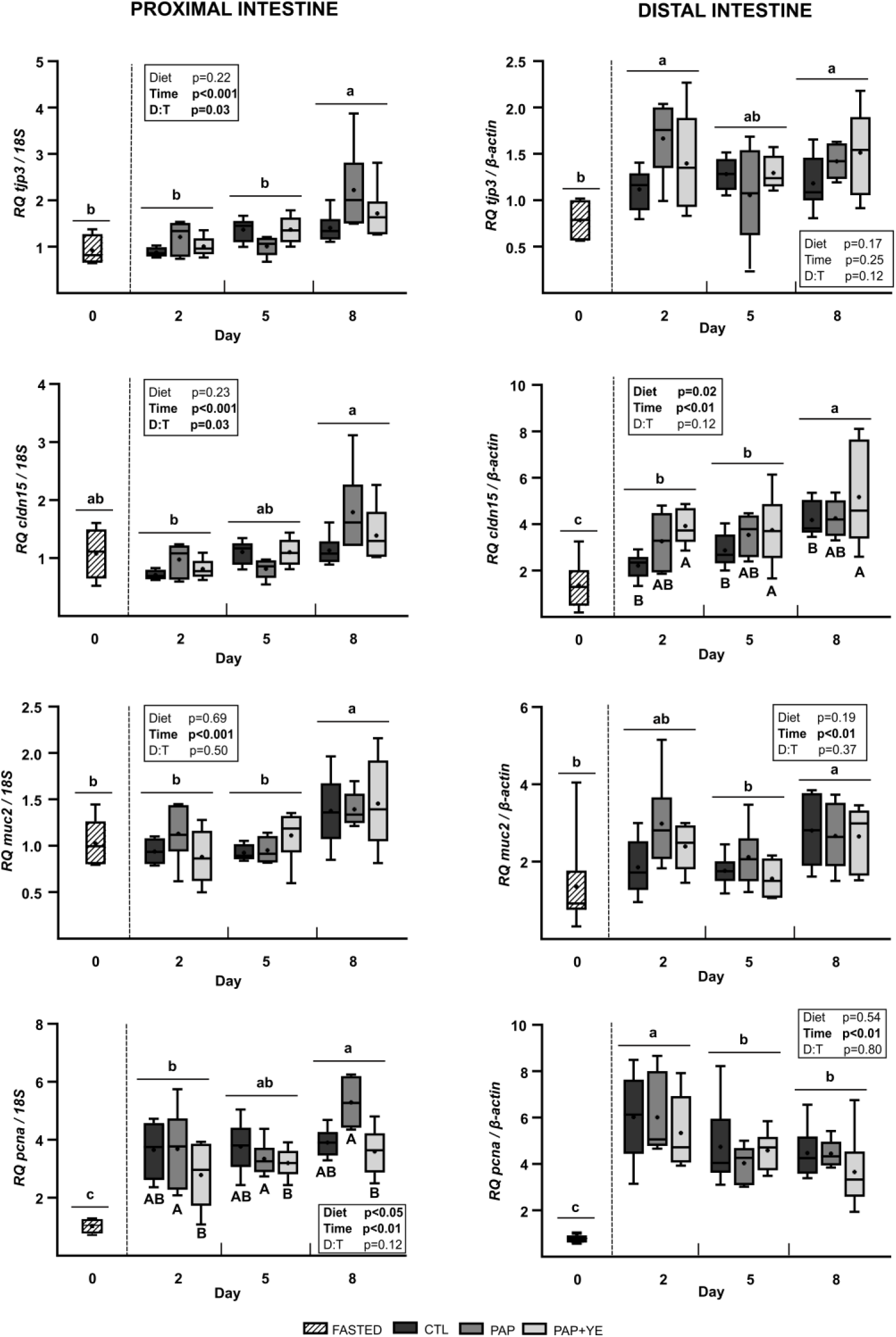
Mean normalized expression (+ standard deviation) (n = 6) of genes related to structure, mucus synthesis and cell proliferation in the proximal and distal parts of the intestine of rainbow trout after fasting and on three of the eight subsequent days of refeeding. CTL, commercial-like feed; PAP, processed animal protein feed; PAP+YE, PAP with 3% yeast extract; *tjp3*, tight junction protein 3; *cldn15*, claudin-15; *muc2*, mucin-2; *pcna*, proliferating cell nuclear antigen. Insets indicate the significance of effects of diet, time and their interaction (D:T). Lowercase letters indicate significant differences between sampling days. Uppercase letters indicate significant differences between diets.

For genes involved in coagulation, expression of the fibrinogen beta chain (*fbg*) gene was too low (Ct ≥ 32) to be quantified adequately, but that of the transcript *cd9* was significantly upregulated in the proximal and distal intestine during refeeding (Fig. 7). In the proximal and distal intestine, *cd9* expression was low in fasted fish and increased after 8 days of refeeding (one-way ANOVA, p < 0.001 and p < 0.05, respectively). However, diet did not influence its expression in either section of the intestine (two-way ANOVA, p > 0.05).

**Figure 7.**
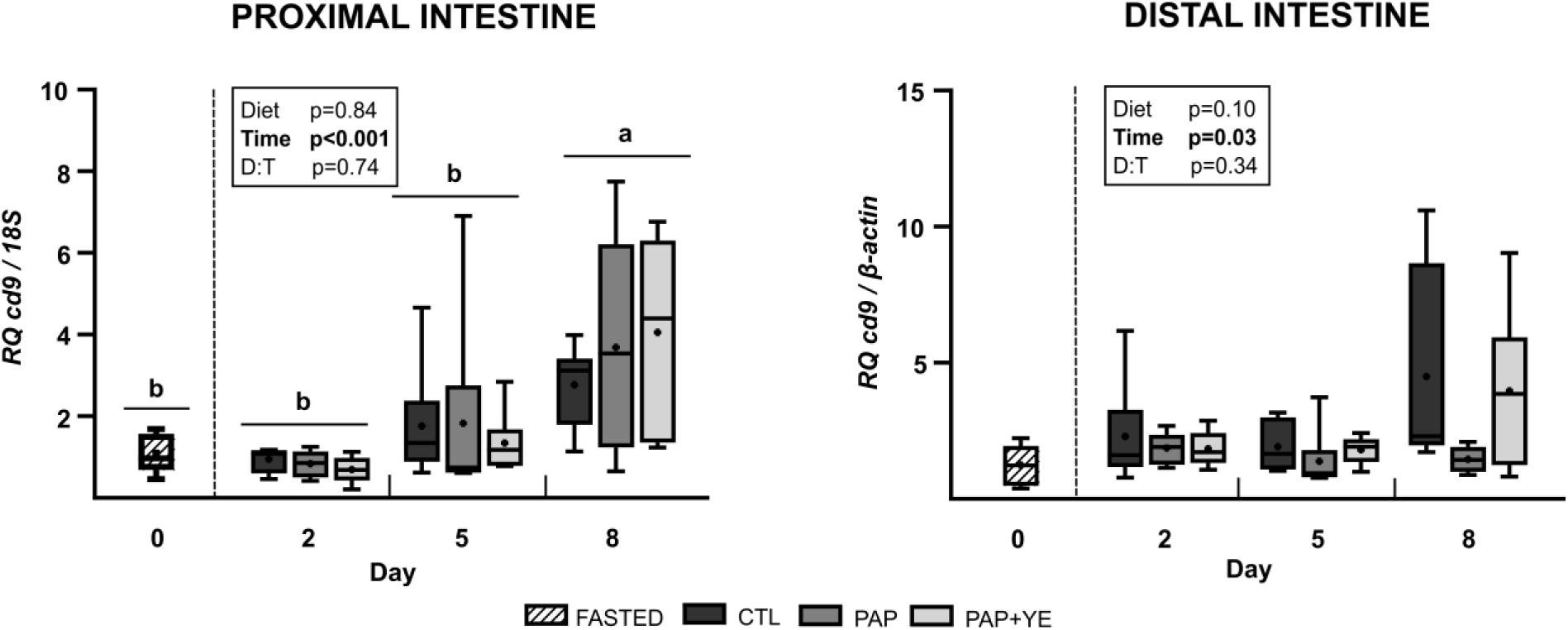
Mean normalized expression (+ standard deviation) (n = 6) of the *cd9* gene, involved in coagulation, in the proximal and distal intestine of rainbow trout after fasting and on three of the eight subsequent days of refeeding. CTL, commercial-like feed; PAP, processed animal protein feed; PAP+YE, PAP with 3% yeast extract; *cd9*, CD9 protein. Insets indicate the significance of effects of diet, time and their interaction (D:T). Lowercase letters indicate significant differences between sampling days.

Expression of selenoprotein S (*selenos*) gene, involved in the endoplasmic reticulum stress response, was strongly down-regulated in fasted fish in both sections of the intestine (one-way ANOVA, p < 0.01) (Fig. 8). It increased on day 2 in refed fish and remained high in the proximal intestine until day 8, while it decreased in the distal intestine without returning to the expression level of fasted fish (two-way ANOVA, p < 0.001). A significant interaction between diet and time was observed in the proximal intestine, indicating strong upregulation of *selenos* expression in fish fed the CTL diet on day 5 and in fish fed the PAP diet on day 8, compared to fish fed the CTL diet on day 8 (two-way ANOVA, p < 0.001).

**Figure 8.**
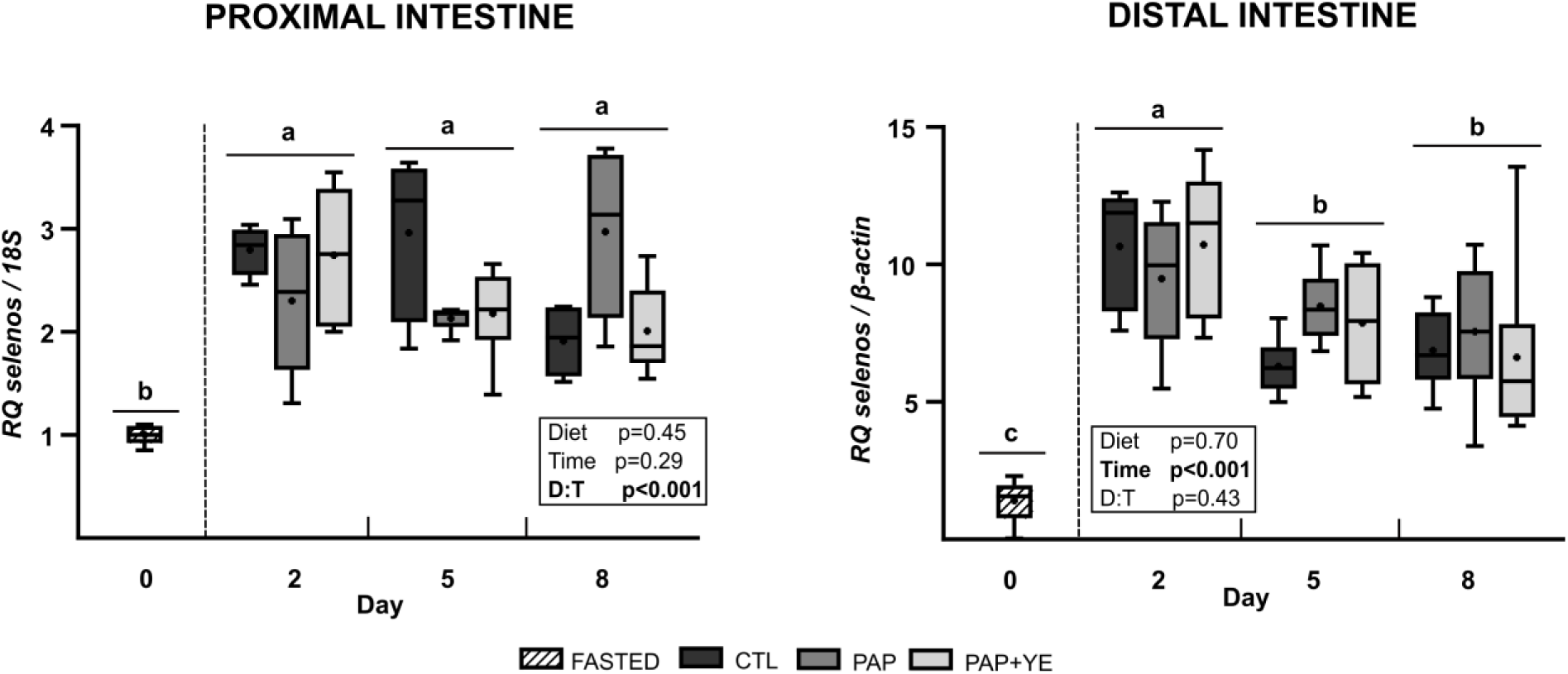
Mean normalized expression (+ standard deviation) (n = 6) of the selenoprotein S (*selenos*) gene, involved in the endoplasmic reticulum stress response, in the proximal and distal intestine of rainbow trout after fasting and on three of the eight subsequent days of refeeding. CTL, commercial-like feed; PAP, processed animal protein feed; PAP+YE, PAP with 3% yeast extract. Insets indicate the significance of effects of diet, time and their interaction. Lowercase letters indicate significant differences between sampling days.

## 3. Discussion

We investigated the early effects of a fishmeal-free diet containing terrestrial animal by-products and yeast-extract supplementation on the structure of the intestine and the inflammatory and immune response of rainbow trout. To avoid potential residual effects of the commercial feed used before and during acclimation, the fish first fasted for four days to empty their digestive tracts. Early responses to the diets were evaluated by quantifying plasma markers of innate immunity and performing histological analysis of the intestinal mucosa. We also investigated the early influence of refeeding on the hepatic and intestinal expression of genes involved in immunity, inflammation, intestinal structure, coagulation and stress response, some of which had been found in a previous study to be significantly influenced by the same diets after 12 weeks of feeding (25).

Feeding diets that contain processed animal proteins and no fishmeal decrease growth performance, but this can be counteracted by adding yeast extract (25). In the long term, it has been shown that these growth performances are accompanied by changes in the transcriptomic profiles of the liver and especially the intestine. The modulated genes were related mainly to the innate immune response, inflammation, coagulation and the response to cellular or pathogenic stress. As previously observed after 12 weeks of feeding (25), the diets did not induce signs of systemic or intestinal inflammation in juvenile rainbow trout. The genes whose expression was modified in the long term were not differentially expressed in the short term, suggesting that the long-term effects were chronic rather than acute, which is the opposite of observations of more sensitive salmonid species, such as Atlantic salmon. The early symptoms of diet-related inflammation in Atlantic salmon are accompanied by an immune response, shriveling of the intestinal villi, increase mucus production by goblet cells and transcriptomic changes (15). Regardless of the diet fed during refeeding, we observed no significant effect on plasma immune markers or the structure of the intestinal mucosa. Of the parameters studied, only two intestinal genes were significantly modulated by the diet: *pcna* and *cldn15*. Over the eight days of refeeding, *pcna* was upregulated in the distal intestine of fish fed the PAP diet, which could indicate and early compensatory effect. Overall, the results suggest that the diets did not cause inflammation, even moderate. Early benefits of including a yeast extract in the diet have been described for common carp (C*yprinus carpio*) (34) and European sea bass (*Dicentrarchus labrax*) (35), but we were not able to demonstrate that doing so had an early influence on the immune response or intestinal structure of rainbow trout. Only the expression of *cldn15* was upregulated in the proximal intestine in response to yeast-extract supplementation, suggesting that it can stimulate intestinal cell adhesion after dietary stress and modulate tissue permeability by preventing the entry of pathogens or toxic molecules (36).

Nonetheless, short-term fasting seemed to induce major changes in the structure of the digestive tract and modulate the physiology of the liver and intestine. Intestinal mucosa and the liver are key organs for metabolism, detoxification and maintenance of homeostasis and play a major role in immunity and protective functions. Although fish can survive fasting for a long period, it soon causes structural changes, induces the response of immune mediators and modulates transcriptomic responses (37–42). During fasting, structural changes include atrophy of the digestive tract and a decrease in the length of the gut, followed by intestinal weight loss and a decrease in the surface area of the intestinal mucosa. A decrease in gut length and mass has been observed in neon damselfish (*Pomacentrus coelestis*) subjected to 16 days of fasting, in javelin goby (*Synechogobius hasta*) on day 7 of fasting (43, 44), and in common carp (*Cyprinus carpi*) subjected to 6 months of starvation (45). As shown for traíra (*Hoplias malabaricus*), a decrease in gut length can also be accompanied by a decrease in the height of intestinal mucosa and the surface area of villi (46). In the present study, short-term fasting had no significant influence on intestinal villi height in the proximal or distal intestine, suggesting that it did not influence mucosa thickness greatly. However, feeding trout after 4 days of fasting significantly increased villi surface area in the proximal intestine on day 8, which indicates a strong relation between refeeding and absorptive capacity. This increase in villi surface area is consistent with upregulation of *pcna,* which has been observed in the intestine of frogs (*Xenopus laevis*) refed after fasting (47). Although this effect was not observed in the distal intestine, our results are consistent with previous observations for salmonids. For short- or medium-term fasting, the gut responds rapidly to fasting, and the effects seem to be reversible upon refeeding, as described for rainbow trout and Atlantic salmon (48, 49).

Genes *tjp3* and *cldn15*, which are involved in enterocytes adhesion and intestinal permeability, were downregulated in the trout during fasting and the first five days of refeeding, and then up-regulated after eight days of refeeding, which confirms the influence of short-term fasting on gut structure. The response of tight junction proteins differed between the sections of the intestine: the distal section seemed to respond more quickly to refeeding, which indicates its higher sensitivity to nutritional status of the fish (50). Like tight junction proteins, goblet cells play an important role in maintaining intestinal homeostasis since they produce intestinal mucus and are present throughout the digestive tract. As a lubricant, mucus facilitates passage of feed nutrients through the digestive tract and protects the intestinal mucosa. During fasting, the number of goblet cells in the intestinal mucosa of teleost fish, such as northern pike (*Esox lucius*) and southern catfish (*Silurus meridionalis*), can decrease significantly to redirect energy to other physiological functions (51–53). Although fasting did not influence the density of goblet cells in the proximal intestine of rainbow trout, two days of refeeding increased it, likely in response to the reactivation by feed in the digestive tract. Conversely, the observed modulation of expression of *muc2* (i.e. low in fasted fish but higher in both sections of the intestine after eight days of refeeding) supports the observations for northern pike and southern catfish. Regarding the phenotype highlighted for intestinal villi surface area and goblet cell density, short-term fasting appeared to influence the proximal intestine more than the distal intestine and refeeding influences goblet cell density and villi surface area significantly.

No significant changes were observed in the activities of the plasma alternative complement pathway, lysozyme, antiprotease or peroxidase. In fish, fasting influences plasma immune markers within the first few days of fasting, and its influence is likely to increase if fasting lasts more than two weeks. The complement pathway, part of innate immunity, groups proteins involved in pathogen lysis and elimination. In fish, its plasma activity can be influenced by short- or long-term fasting. For example, tinfoil barb (*Barbonymus schwanenfeldii*) and crucian carp (*Carassius auratus*), exposed to short- (two weeks) and long-term fasting (60 days), respectively, showed a significant decrease in plasma alternative complement activity. Thus, nutrient deprivation could cause the innate immune system to fail, which could decrease the ability to fight pathogens, but this was not observed in rainbow trout, likely due to the short fasting period (54, 55). Lysozyme, also part of innate immunity, protects against bacterial infections when released by phagocytes. In the present study, the short fasting period did not seem to influence plasma lysozyme activity in rainbow trout, as it was similar for fasted and refed fish. This agrees with the literature, which indicates that the influence of fasting on plasma lysozyme activity depends greatly on the species studied, its physiological status and the duration of fasting (56). For tinfoil barb, Eslamloo et al. (54) showed that one week of fasting was sufficient to induce an increase in lysozyme activity, which then decreased sharply after two weeks of fasting. In Chinese sturgeon (*Acipenser sinensis*), 43 days of fasting clearly induced lysozyme activity (57), while prolonged fasting did not influence it in European sea bass, red seabream (*Pagellus bogaraveo*) or European eel (*Anguilla anguilla*) (58). Peroxidase is an antioxidant enzyme involved in eliminating reactive oxygen species released during oxidative stresses that are induced, for example, in Chinese perch (*Siniperca chuatsi*) by fasting (59). Several studies of fish show that plasma peroxidase can be induced soon after fasting begins. For example, plasma peroxidase activity in binni (*Mesopotamichthys sharpeyi*) fingerlings was strongly induced on day 8 of fasting (60), and gilthead seabream (*Saprus aurata*) had the highest peroxidase activity after 7 days of fasting (61). However, some studies have shown that if fasting exceeds several weeks, peroxidase activity may sharply decrease due to fatigue of the antioxidant system (54, 62). In the present study, peroxidase activity was similar for fasted and refed fish, suggesting that the four days of fasting may have been too short to increase peroxidase activity in rainbow trout. Plasma antiproteases protect against pathogens by inhibiting the released proteases. Little is known about the influence of fasting on antiprotease activity, and existing studies focus mainly on tissues other than blood or analyze antiprotease activity after a bacterial challenge (54, 63, 64). However, in a study of Nile tilapia (*Oreochromis niloticus*) fasted for 21 days, plasma antiprotease activity significantly increased but then decreased during refeeding. Rather than being immune-related, these effects were due to inhibiting proteases that degrade dietary proteins and thus prevent them from being digested and assimilated during prolonged fasting (65). Overall, our results suggest that the fasting period was too short to induce a systemic immune response or protein catabolism in rainbow trout.

Although we did not observe systemic or histological inflammation in response to the four-day fast, the expression of certain genes related to immunity, inflammation and cell protection in the liver and intestine differed between fasted and refed fish. In agreement with observations on red porgy (*Pagrus pagrus*), Atlantic salmon (*Salmo salar*) and Japanese grenadier anchovy (*Coilia nasus*) (66–68), expression of the complement *c3* gene in rainbow trout was higher after two days of refeeding, although it was not significantly lower in fasted fish, perhaps because the fasting period was shorter than those in previous studies. The protein encoded by the *a2m* gene was upregulated in trout subjected to a four-day fast, suggesting early regulation of acute phase protein homeostasis after short-term fasting. The opposite was observed in red porgy and Japanese grenadier anchovy, but these fish were subjected to a longer fast, suggesting that prolonged fasting may decrease the activity of this acute phase protein in fish (66, 68). Expression of *prg4*, which encodes a proteoglycan stimulating macrophages during inflammation (69), decreased during refeeding but did not necessarily differ from that of fasted trout. To our knowledge, no study has described the influence of fasting on *prg4* expression in the liver of fish. However, the results obtained for rainbow trout suggest that nutrients must be present to maintain sufficient immune function.

In the intestine, most genes related to the immune and inflammatory responses were not expressed, or their expression was too low to be quantified adequately. Although *ch25h* has been implicated in inflammatory and viral pathologies (70, 71), its high expression in the distal intestine of fasted fish was not sufficient to indicate that it is related to the immune or inflammatory response in the intestine, since the trout had no clear sign of inflammation caused by fasting. However, *il1β* expression strongly but temporarily increased in response to refeeding. Response related to inflammation in the gut seem to occur more frequently when long-term fasting exceeds several weeks (72–75).

Although we could not quantify all genes related to coagulation in the liver and intestine due to low expression, the response of the *cd9* gene in the intestine suggests that fasting decreases wound healing and clotting capacities and that refeeding tends to restore these functions, which agrees with previous studies. Fasting impairs hemostasis and clotting, which may be restored upon refeeding, as observe for traíra (76) and for rainbow trout, in which all transcripts that encoded for proteins involved in blood coagulation were downregulated in the liver during fasting (77).

The *miox* gene, which encodes myo-inositol oxygenase, was upregulated in the liver of fasted rainbow trout compared to fish refed on day 8. In Nile tilapia, myo-inositol is used as an osmolyte to protect cells and maintain normal cytoplasmic osmolarity under hypertonic conditions (78, 79). In plants, specifically thale cress (*Arabidopsis thaliana*), *miox* is upregulated under low nutrients conditions (80). Since fasting can stress cells and impair osmolarity of rainbow trout (77), its upregulation in the liver of fasted fish seems consistent with its protective function. Moreover, its downregulation after eight days refeeding seems to be related to a return to normal osmolarity after refeeding and the presence of nutrients in the gut. In the intestine, especially in the distal section, *selenos* expression, a gene encoding a protein involved in protein folding in the endoplasmic reticulum and in glucose metabolism, was also strongly influenced by the transition from fasting to refeeding. These results agree with those of previous studies and suggest that *selenos* is regulated by glucose deprivation, and that fasting is likely to cause endoplasmic reticulum stress, and consequently a failure of protein folding (81–83).

In conclusion, we were unable to demonstrate early benefits or repercussions associated with feeding rainbow trout a diet that contained animal by-products but no fishmeal on their immune response and gut structure, whether the diet was supplemented with yeast extract or not. Thus, the decrease in growth performances associated with the PAP diet, the improvement of this diet with the addition of yeast extract and the changes in the hepatic and intestinal transcriptomic profile were not related to an early reaction of the fish, but to a long-term adaptation (25). Conversely, short-term fasting seems to have much more influence on the physiology of the liver and intestine. Four days of fasting followed by refeeding significantly increased the intestinal surface and modulated the expression of genes involved in inflammation and immunity, the pathogen response and cellular stress in liver and intestine, with no clear evidence of systemic or tissue inflammation. This difference may be associated with the relatively short duration that rainbow trout were fasted. However, these findings are consistent with the current use of fasting periods on fish farms to increase survival, maintain growth and improve the immune defenses of fish in stressful or epizootic situations (84, 85).

## Acknowledgement

We dedicate this study to Frank Sandres, who cared for the fish during this study and sadly passed away. We are also grateful to the research unit ToxAlim (INRAE, Toulouse, France) for providing their knowledge and lending us the equipment used to perform histology in this study.

## 6. Author contributions

Conceived and designed research: LF, SS, FT

Performed experiments: LF, DP, JB, FT

Analyzed data: LF

Interpreted results: LF

Prepared figures: LF

Drafted manuscript: LF

Edited and revised manuscript: LF, DP, FT, BC, JB, CC, NR, KP, SS

Approved final version of manuscript: LF, DP, FT, BC, JB, CC, NR, KP, SS

